# Identification of genomic regions of dry bean (*Phaseolus vulgaris* L.) associated with agronomic and physiological traits under drought stressed and well-watered conditions using genome-wide association study

**DOI:** 10.1101/2022.11.18.517065

**Authors:** Bruce Mutari, Julia Sibiya, Admire Shayanowako, Charity Chidzanga, Prince M. Matova, Edmore Gasura

## Abstract

Understanding the genetic basis of traits of economic importance under drought stress (DS) and well-watered (NS) conditions is important in enhancing genetic gains in dry beans (*Phaseolus vulgaris* L.). This research aims to: (i) identify markers associated with agronomic and physiological traits for drought tolerance and (ii) identify drought-related putative candidate genes within the mapped genomic regions. An Andean and Mesoamerican diversity panel (AMDP) comprising of 185 genotypes was screened in the field under drought stress (DS) and well-watered (NS) conditions for two successive seasons. Agronomic and physiological traits, *viz*., days to 50% flowering (DFW), plant height (PH), days to physiological maturity (DPM), grain yield (GYD), 100-seed weight (SW), leaf temperature (LT), leaf chlorophyll content (LCC) and stomatal conductance (SC) were phenotyped. Principal component and association analysis were conducted using filtered 9370 Diversity Arrays Technology sequencing (DArTseq) markers. The mean PH, GYD, SW, DPM, LCC and SC of the AMDP was reduced by 12.1, 29.6, 10.3, 12.6, 28.5 and 62.0%, respectively under DS. Population structure analysis revealed two sub-populations, which correspond to the Andean and Mesoamerican gene pools. Markers explained 0.08 – 0.10, 0.22 – 0.23, 0.29 – 0.32, 0.43 – 0.44, 0.65 – 0.66 and 0.69 – 0.70 of the total phenotypic variability (*R*^*2*^) for SC, LT, PH, GYD, SW and DFW, respectively under DS conditions. For NS, *R*^*2*^ varied from 0.08 (LT) to 0.70 (DPM). Overall, 68 significant (p < 10^−03^) marker-trait associations (MTAs) and 22 putative candidate genes were identified across DS and NS conditions. Most of the identified genes had known biological functions related to regulating the response to moisture stress. The findings provide new insights into the genetic architecture of moisture stress tolerance in common bean. The findings also provide potential candidate SNPs and putative genes that can be utilized in gene discovery and marker-assisted breeding for drought tolerance after validation.

## Introduction

Common bean (*Phaseolus vulgaris* L., 2*n* = 2x = 22) is one of the major pulse crops consumed worldwide with a relatively small diploid genome size of approximately 473 Mb [1]. It is a cheap source of proteins and important micronutrients such as iron (Fe) and zinc (Zn) for millions in many African and Latin American countries [2, 3]. Beebe et al. [4] reported that Sub-Saharan Africa (SSA) and Latin America produce the largest volume of common beans, representing more than 60% of the world’s bean production. Common bean was subjected to two parallel domestication events on the American continent, resulting in two different primary gene pools namely the Andean and the Mesoamerican [5, 6]. The Andean gene pool originated from the Andes mountains of South America and consists of medium (25 - 40 g per 100 seeds) or large (≥ 40 g per 100 seeds) seeded genotypes [7]. On the other hand, the Mesoamerican gene pool is native to Central America and Mexico, and comprises of small seeded genotypes (≤ 25 g per 100 seeds). According to Bitocchi et al. [8], there is more genetic variation within the Mesoamerican gene pool compared to the Andean gene pool.

Common beans are notably sensitive to climatic and environmental variations. This is aggravated by the fact that most bean growing regions in the world experience different production constraints including intermittent and terminal drought stress which adversely affect grain yield [9–12]. As reported by Katungi et al. [13], 73% of common bean production in SSA occurs in environments which experience moderate to severe drought stress. Beebe et al. [4], Hoyos-Villegas et al. [14] and Valdisser et al. [15] reiterated that drought stress is the most important grain yield-limiting abiotic factor of dry bean worldwide. It is predicted from various climate models that the duration and frequency of droughts are expected to increase in SSA [16]. Drought stress reduces stomatal conductance, total chlorophyll content, leaf expansion, number of days to physiological maturity, seed yield and biomass, number of pods and seeds per plant, seed size and harvest index [17–22]. According to Asfaw et al. [23], severe drought stress can result in grain yield losses of up to 80%. In Zimbabwe, grain yield reductions of more than 50% were reported by Mutari et al. [24] under terminal drought stress.

As reported by Mutari et al. [25], bean farmers in Zimbabwe have been using different mitigation strategies to minimize grain yield losses due to terminal drought stress. These strategies include soil mulching, ridging, cultivating the soil to retain more moisture and reducing the area under the bean crop. However, host plant resistance is a more sustainable, environmentally friendly and labour saving technology for managing drought stress in common beans compared to the multiple cultural practices. For this reason, most dry bean breeding programmes aim to introduce drought tolerance into new cultivars to address the needs and preferences of smallholder farmers in the face of climate change [26].

Several researchers have successfully used different types of deoxyribonucleic acid (DNA)-based marker systems in association mapping of complex traits in common beans. The most widely used marker systems include simple sequence repeats (SSRs; [27–29]), amplified fragment length polymorphisms (AFLPs; [28, 30]), single nucleotide polymorphisms (SNPs; [3, 14, 31–34]) and microarray based Diversity Arrays Technology (DArT; [15, 35]) markers. However, SNP markers are widely preferred in marker assisted selection (MAS), genetic diversity analyses, genomic selection, haplotype mapping, genome wide association studies (GWAS), linkage map construction and population genetics [36]. They are widely preferred because they exhibit high level of polymorphism and occur in abundance (cover the whole genome) as differences of individual nucleotides between individuals.

Understanding the underlying genetic architecture of agronomic and physiological traits under drought stress (DS) and well-watered (NS) conditions is a fundamental prerequisite for the genetic improvement of these traits in common beans using MAS. Thus, dissecting the genetic basis of multiple polygenic traits of economic importance such as drought tolerance with respect to the genomic regions and/or genes involved and their effects is important to improve genetic gains in breeding for superior grain yield in dry beans under DS and NS environments. This can be accomplished through complementary approaches such as GWAS and genomic prediction models [6]. Genome wide association study is a powerful tool for characterizing the genetic basis of quantitative traits, and identifying multiple candidate genes (marker alleles) associated with variation in quantitative traits (marker-trait associations; MTA) of interest in crop species using high density DNA markers at high level of genetic resolution [34, 37–41].

Genome wide association study is also known as association mapping (AM) or linkage disequilibrium (LD) mapping [42]. It is based on linkage LD and historical recombination events of alleles of detected quantitative trait loci (QTL) at relatively high level of genetic resolution due to high genetic variability in the diverse population such as landraces, elite breeding lines and improved cultivars [43, 44]. The historical recombination events would have naturally occurred during the evolution and domestication of the crop, and crop improvement (several generations) [33]. With GWAS, the mapping resolution is increased as a result of the high number of recombination events in the genetically diverse genotypes within the natural population [45]. Therefore, it is inexpensive and reduces research time (no need to develop a mapping population) with greater allele numbers. The identification of genomic regions and diagnostic genetic markers associated with grain yield and yield-attributing traits under DS and NS conditions will facilitate trait introgression and marker assisted selection (MAS).

Genome wide association study has been successfully used to detect MTAs and QTLs in common bean. Several QTLs associated with disease and insect pest tolerance have been identified in dry bean [32, 46–50]. Similarly, MTAs were identified for drought tolerance traits in dry bean [14, 15, 51–53]. Also, MTAs were identified for nutritional composition-related traits [6, 33], symbiotic nitrogen fixation [54], cooking time [55] and photosynthetic traits [34, 56] in dry bean. Genomic regions governing agronomic traits in DS and yield potential environments were also identified in dry bean [1, 6, 14, 34, 57]. Even though several significant MTAs were identified in previous GWAS studies for agronomic traits in DS environments, the use of very low thresholds (-log_10_ *p*-value ≥ 3.0) in most of the studies in determining significant MTAs might have resulted in many false positives. In addition, despite the fact that several QTLs/MTAs associated with agronomic traits have been identified in dry bean, further genetic studies are required using different genetic backgrounds to reach a saturation point. Moreover, most of the reported putative genes for agronomic and physiological traits were detected under yield potential environments.

Additionally, some of the previous mapping studies [14, 17, 51, 58–61] conducted on agronomic and physiological traits used a small population size and a limited number of molecular markers. This resulted in QTL with low resolution or poor estimation of marker effects, making it difficult to make inferences on putative candidate genes correlated with the identified QTL. Moreover, some of the previously identified QTLs explained low total genetic variance [23], and were sometimes not stable across environments due to genotype by environment interaction (GEI) [52]. Thus their potential for MAS in developing genotypes that are tolerant to drought stress was inconclusive. Therefore, additional studies are required to dissect the genetic basis of agronomic and physiological traits in dry bean under DS and optimal environments for increased genetic gains. The objectives of this study were: (i) to identify single nucleotide polymorphism (SNP) markers significantly associated with agronomic and physiological traits for drought tolerance and; (ii) to identify drought-related putative candidate genes associated with traits within the mapped genomic regions.

## Materials and Methods

### Description of the study location

The field experiments (drought stress; DS and well-watered; NS) were conducted at the screening site for moisture stress tolerance located at Save Valley Experiment Station (SVES), Zimbabwe. The experiments were carried out during the 2019 and 2020 dry winter seasons (April – July). Save Valley Experiment Station is characterised by clay soils and is located in the drier lowveld region of Zimbabwe where dry beans are commercially produced during the dry winter season (Table 1). The research station receives an average annual rainfall of 450 mm that is usually distributed between the months of December and April. In both seasons, no precipitation was received during the trial evaluation period. Historically, SVES presents few rainfall occurrences during the dry winter season [24]. Daily temperatures (°C) and relative humidity (%) were recorded with a digital weather station (Table 1) during the growing seasons. More details on the agro-ecological characteristics of SVES are outlined in Table 1.

**Table 1.** Geographic information system, monthly weather conditions and soil characteristics during the growing seasons at Save Valley Experiment Station, Zimbabwe (April to July, 2019 and 2020).

| Parameter | 2019 dry season |  |  |  |  | 2020 dry season |  |  |  |
| --- | --- | --- | --- | --- | --- | --- | --- | --- | --- |
|  |  | April | May | June | July | April | May | June | July |
| Temperature (°C) | Max | 33.00 | 29.00 | 28.00 | 30.00 | 31.00 | 28.50 | 27.00 | 32.00 |
|  | Min | 9.00 | 9.50 | 10.00 | 12.00 | 11.50 | 8.00 | 8.5.00 | 12.50 |
| Relative Humidity (%) | Max | 82.00 | 95.00 | 69.00 | 91.00 | 74.00 | 85.00 | 69.00 | 71.00 |
|  | Min | 42.00 | 56.00 | 44.00 | 25.00 | 46.00 | 59.00 | 50.00 | 30.00 |
| Total Rainfall (mm) |  | 0.00 | 0.00 | 0.00 | 0.00 | 0.00 | 0.00 | 0.00 | 0.00 |
| Soil type |  | Clay |  |  |  | Clay |  |  |  |
| Latitude |  | 20°32'S |  |  |  | 20°43'S |  |  |  |
| Longitude |  | 33°09'E |  |  |  | 33°03'E |  |  |  |
| Altitude (m.a.s.l) |  | 452 |  |  |  | 449 |  |  |  |

masl = meters above sea level, mm = millimetres, ppm = parts per million, Max = maximum and Min = minimum.

**Table 1.**
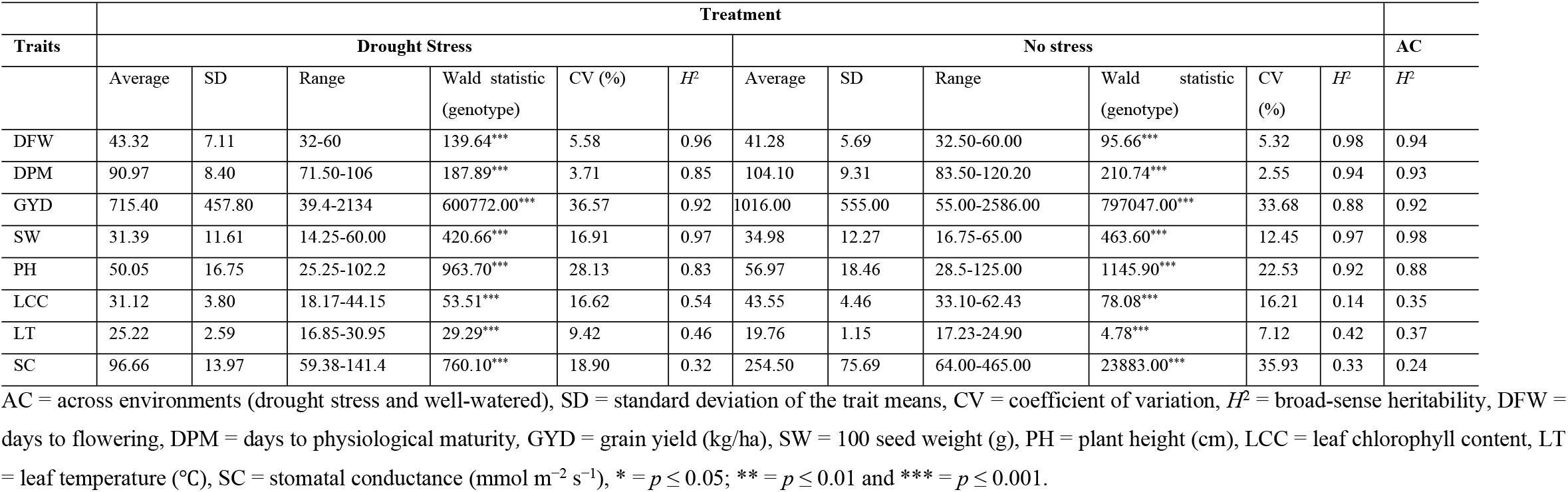
Phenotypic summary statistics, coefficient of variation and broad-sense heritability of the measured traits for all the 185 dry bean genotypes based on the best liner unbiased prediction (BLUP) value grown under drought stressed and non-stressed conditions.

### Germplasm

A total of 185 dry bean genotypes constituted the Andean and Mesoamerican diversity panel (AMDP). The AMDP comprised of landrace collections (25), released cultivars (18) and elite breeding lines (142) of different market classes such as sugars, calimas, small whites, large whites and large red kidneys (S1 Table). The genotypes were sourced from public and private breeding institutions located in different geographic regions. These included the Alliance of Bioversity International and International Center for Tropical Agriculture (ABC) in Colombia (87), ABC in Malawi (67), ABC in Uganda (18), Ethiopian Institute of Agricultural Research (EIAR) in Ethiopia (3), Crop Breeding Institute in Zimbabwe (6) and Seed-Co, also in Zimbabwe (4) (S1 Table).

### Field phenotyping of the diversity panel

#### Experimental design, irrigation scheduling and trial management

The AMDP was evaluated side by side under DS and NS treatment conditions during the 2019 and 2020 dry winter seasons. In both seasons, the genotypes in both DS and NS treatments were established in a 5 × 37 alpha lattice design with two replications. The seepage of water from the NS treatment to the DS treatment was minimized by maintaining a 30 m buffer zone between the two treatments. Each genotype was hand planted in four-row plots of 3 m in length, and an inter-row spacing of 0.45 m. Compound D (N = 7%, P = 14%, K = 7%) was applied at planting at a rate of 300 kg/ha. Ammonium nitrate (34.5% N) was applied in both DS and NS treatments as a top-dressing fertilizer thirty days after emergence at a rate of 100 kg/ha. An overhead sprinkler irrigation system was used to irrigate both DS and NS treatments during both seasons of evaluation. The irrigation cycles in both DS and NS treatments were as described by Mutari et al. [24]. In both seasons, recommended agronomic practices were followed for the management and control of pests such as diseases, insects and weeds.

#### Collection of data on agronomic and physiological traits

At the flowering stage of growth, the number of days from planting to 50% flowering (DFW) were recorded in both treatments. The DFW was recorded when 50% of the plants in a plot had at least one or more open flowers. At mid-pod filling, leaf temperature (LT; °C), stomatal conductance (SC; mmol m^-2^ s^-1^) and leaf chlorophyll (LCC) content were collected on all genotypes in both DS and NS treatments. The LT and SC data were recorded from the surface of the uppermost fully expanded young leaf between 11:00 am to 14:00 pm using a FLUKE precision infrared thermometer (Everest Interscience, Tucson, AZ, USA) and a hand-held leaf porometer (Decagon Devices®, Pullman, WA, USA), respectively. Three readings were collected on three different randomly chosen plants from each plot per replicate in both the DS and NS treatments. The three measurements were averaged to obtain one final reading per plot. Phenotyping for LT and SC was done for an average of six days on clear, sunny days with minimal wind. Regarding the LCC, this was measured using a soil and plant analysis development (SPAD) chlorophyll meter (SPAD-502*Plus*, Konica-Minolta, Osaka, Japan) on two fully developed leaves of three plants in each plot. Then, the average value was calculated. At physiological maturity, the following traits were recorded from the two inner rows from every plot for every genotype in both treatments and seasons: plant height (PH; cm), days from planting to physiological maturity (DPM), grain yield (GYD; kg/ha) and 100-seed weight (SW; g). Plant height which was measured from the base of the plant (soil surface) to the top node bearing at least one dry pod with seed was averaged from three plants per plot. The DPM were recorded as the average number of days from planting to when 95% of pods in a plot lose their green colour. Grain yield was recorded from the two middle rows in each plot using a weighing scale, and converted to kilograms per hectare (kg/ha) at 12.5% moisture basis. The SW was determined using a beam balance weighing scale by measuring the weight of 100 seeds randomly from each plot harvest.

### Statistical analysis of phenotypic data

Before conducting analysis of variance, normality tests were conducted in Genstat® Discovery 18th Edition [62] using residuals of the agronomic and physiological traits. The agronomic and physiological traits were analysed in Genstat® Discovery 18th Edition [62] using mixed models from which the best linear unbiased predictors (BLUPs) were obtained. The BLUPs were estimated for the studied traits to minimize the environmental and seasonal effects. The BLUPs for each entry were estimated through individual environment (DS or NS) analysis, and by combined analysis (across water regimes). In the first step of analysis (single-environment analysis), the phenotypic data of each individual environment were analysed using a mixed linear model (MLM). In this model, blocks and genotypes were treated as random effects, and replications were considered as fixed effects. Genotype effects were declared to be random to enable the calculation of BLUPs and broad-sense heritability (*H*^*2*^). The MLM presented below was fitted:

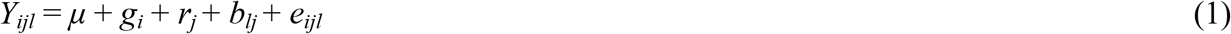

where *Y*_*jkl*_ = is the phenotypic observation of the genotype *i* in replicate *j* in block *l* within replicate *j, μ* = grand mean effect, *g*_*i*_ = random effect associated with genotype *i, r*_*j*_ = fixed effect associated with replicate *j, b*_*lj*_ = random effect associated with block *l* nested within replicate *j*, and e_ijl_ = residual effect associated with observation *ijl*. For a combined or multi-environment analysis, a MLM was used. In this model, blocks nested within replications, replicates nested within environments, genotypes and their interactions with environments (GEI) were considered as random effects. Environments, defined as year × water regime combination were considered as fixed effects. The MLM presented below was fitted:

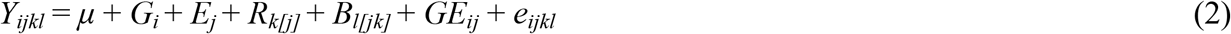

where *Y*_*ijkl*_ = effect of genotype *i* in environment *j* and *k*th replication within environment *j* and *I*th block nested within replicate *k* and environment *j, μ* = grand mean, *G*_*i*_ = random effect of the *i*th genotype, *E*_*j*_ = fixed effect of the *j*th environment, *R*_*k[j]*_ = random effect associated with the replicate *k* nested within environment *j, B*_*l[jk]*_ = random effect of block *l* nested within environment *j* and replicate *k, GE*_*ij*_ = random effect of the interaction between genotype *i* and environment *j*, and *e*_*ijkl*_ = random error associated with observation *ijkl*. The analysis was performed using the Restricted Maximum Likelihood (REML) method implemented in GenStat 18th edition [62]. Broad-sense heritability estimates for the agronomic and physiological traits were calculated following the formula proposed by Cullis et al. [63]. Heritability was classified as low when less than 30 %, moderate when between 30-60 % and high when more 60 % [64]. Drought intensity index (DII) at the location, percentage GYD reduction (%GYR) due to DS, drought susceptibility index (DSI), geometric mean productivity (GMP) and drought tolerance index (DTI) of each entry were calculated as described by Mutari et al. [24]. A ranking method was used to select superior drought tolerant genotypes by calculating the mean rank of each genotype across all the studied indices.

### Genotyping of the diversity panel

Genomic DNA of the 185 genotypes was extracted from young leaves of 2-week old bean plants following the plant deoxyribonucleic acid (DNA) extraction protocol for Diversity Arrays Technology (DArT; [65]). A NanoDrop Spectrophotometer (ND-8000, NanoDrop Technologies, Inc.) was used to determine the concentration of the DNA. The agarose gel (1% agarose gel) electrophoresis was used to evaluate the quality of the DNA. The DNA from the samples used in this study were genotyped using the Diversity Arrays Technology Sequencing (DArTseq) protocol using a set of 24,450 silico DArT markers. The DArT markers used were evenly distributed across all 11 chromosomes of common bean. Genotyping by sequencing (GBS) was done at the Biosciences Eastern and Central Africa (BecA) Hub of the International Livestock Research Institute (BecA-ILRI) in Kenya. The silico DArTs used had polymorphic information content (PIC) values ranging from 0.01 to 0.50, reproducibility values of 1.00, and the proportion of missing data per marker was 7% (mean call rate of 93%, ranging from 81 to 100%). The entire data set of SNP markers was filtered in TASSEL v5.2 [66] to remove SNP loci with unknown physical positions on the common bean genome, monomorphic SNPs, and SNP markers with more than 20% missing data and minor allele frequency (MAF) of less than 5% (<0.05) threshold [15, 49, 67]. A final total of 9370 (38%) DArTseq SNPs distributed across the 11 chromosomes were retained after filtering for use in association analysis and population structure analysis via principal component analysis (PCA).

### Inference of population structure

The genotypic data was imputed for missing alleles of SNPs on the KDCompute online sever (https://kdcompute.igs-africa.org/kdcompute/) using the optimal imputation algorithm to increase the power of the study. KDCompute was also used to graphically visualize the distribution of SNPs across the common bean genome. The population genetic structure was determined based on the Bayesian model-based clustering approach using the Bayesian inference program in STRUCTURE software version 2.3.4 [68]. A subset of additionally filtered SNP markers (4095) at or near Hardy-Weinberg equilibrium (r^2^ < 0.8) and that covered the entire genome were used in population structure analysis with STRUCTURE [14, 15, 31]. This was done to reduce the background and admixture linkage disequilibrium (LD) owing to linked loci [68].

Settings for the STRUCTURE program were set as follows to derive the population structure: a burn-in period length of 10,000, and after burn-in, 10,000 Markov Chain-Monte Carlo (MCMC) repetitions. The number of sub-populations or clusters (K) was set from 1 to 10, with ten independent runs for each *K* [3, 48, 55]. The best K-value explaining the population structure was inferred using the Delta *K* (ΔK) method in Evanno et al. [69] implemented in the on-line tool structure harvester software [70]. Genotypes with ancestry probability/coefficient ≥ 0.90 (≥ 90%) (pure genotypes) for the Andean sub-population were allocated to the Andean gene pool [31, 71] (S1 Table). On the other hand, genotypes with ancestry probability ≥ 0.90 for the Mesoamerican sub-population were allocated to the Mesoamerican gene pool. Those with ancestry probability < 0.90 were considered as admixed [71]. The clustering of the AMDP was further assessed and visualized in a 3D scatter plot using PCA in prcomp R 3.0 function [72].

### Marker-trait association tests and linkage disequilibrium analyses

The filtered 9370 SNPs and the adjusted trait means (BLUPs) for each of the environments (DS and NS) were used as input data in marker-trait association (MTA) analysis. The more conservative compressed mixed linear model (CMLM) procedure in the genome association and prediction integrated tool (GAPIT) (v3) program of R software was used to determine the MTAs following the *Q* + *K* model according to Lipka et al. [73]. *Phaseolus vulgaris* is characterised by a strong genetic structure necessitating the need to use the Q + K model [74]. The CMLM incorporated both the population structure (*Q*; fixed effect) and kinship (*K*; random effect) matrices as covariates to correct the population structure, increase statistical power of the analysis and minimize false positives (spurious MTAs) [67, 72, 75]. The K matrix was included in the association analysis to correct for cryptic relatedness within the AMDP [54, 67]. The threshold for significant MTA was set at p < 0.001 to reduce the risk of false MTAs.

The Manhattan plots drawn using the CMplot package in R 3.5.3 were used to visualise the significant MTAs for each environment. The p*-*values were plotted as –log_10_(p) to generate the Quantile-Quantile (Q-Q) and Manhattan plots using the CMplot package in R package [76]. The Q-Q plots were produced from the observed and expected logarithm of the odds (LOD) scores for each trait. The LD Heatmap package in R 3.0 was used to generate the LD Heatmaps for the significant markers of each trait [77, 78]. Alleles with positive additive effects resulting in higher values of GYD, SZ and LCC were described as “superior alleles” under both DS and NS conditions, whereas alleles resulting in decreased GYD, SZ, and LCC were “inferior alleles”. On the other hand, alleles with negative effects resulting in lower values of DFW, DPM, LT and SC were considered to be “superior alleles” under DS conditions. The Jbrowse feature on Phytozome v13 was used to browse the *P. vulgaris* G19833 v2.1 reference genome sequence [1] to gain insight into potential putative candidate genes associated with significant SNPs for each trait. The functional annotation of the gene was checked on Phytozome v13 website (http://phytozome.net) to postulate the role of the gene in the control of a target trait.

### Putative candidate gene prediction

Plausible candidate genes were identified based on the window size of 200 kb (maximum ± 100 kb) on either side (upstream and downstream) of the significant marker [74, 79]. The window size of 200 kb is the average LD [74, 79]. A gene was considered a potential candidate using the following criteria: (i) if the gene contained a significant SNP or the gene contained a SNP that was in LD with a significant SNP [3], and (ii) if the gene had a known role related to regulating moisture stress response and plant growth and development under water deficit based on gene ontology term descriptions in Phytozome v13. For the positional candidate genes that did not have adequate functional annotation information on Phytozome v13, the sequence data of the significant SNP was used against NCBI database using the basic local alignment search tool for nucleotide (BLASTn; https://blast.ncbi.nlm.nih.gov/smartblast/smartBlast.cgi).

## Results

### Variations of agronomic and physiological traits under two water regimes

The descriptive statistics and *H*^*2*^ estimates for the agronomic and physiological traits under DS and NS environments are shown in Table 2. Residual maximum likelihood analysis revealed highly significant (p < 0.001) genotypic main effects on all the studied traits under both DS and NS environments supporting the use of the AMDP for GWAS purposes. Overall, phenotypic variability was observed among the genotypes for DFW, LCC, LT, SC, PH, DPM, GYD and SW under DS and NS conditions. High *H*^*2*^ estimates (0.83 - 0.97) were observed for all the studied traits under DS, except for SC (*H*^*2*^ = 0.32), LT (*H*^*2*^ = 0.46), and LCC (*H*^*2*^ = 0.54). Under NS conditions, high *H*^*2*^ estimates (0.88 – 0.98) were observed for all the traits except for LCC (*H*^*2*^ = 0.14), SC (*H*^*2*^ = 0.33), and LT (*H*^*2*^ = 0.42).

**Table 2.** Drought tolerance indices and predicted genotype values for grain yield (across environments) of top 20 drought tolerant genotypes.

| Genotype | Gene pool | GYD (kg/ha) | DSI | GMP | DTI | %GYR | Mean rank |
| --- | --- | --- | --- | --- | --- | --- | --- |
| G184 | Andean | 2222.7 | 0.16 | 2205.3 | 4.74 | 4.69 | 25.5 |
| G176 | Andean | 2097.5 | 0.35 | 2084.0 | 4.25 | 10.60 | 29.9 |
| G147 | Andean | 2080.1 | 1.26 | 1995.6 | 4.03 | 37.83 | 66.3 |
| G146 | Andean | 2067.4 | 1.18 | 1994.5 | 4.02 | 35.49 | 61.5 |
| G158 | Admixed | 2017.1 | -0.01 | 1979.5 | 3.82 | -0.26 | 24.5 |
| G135 | Andean | 1968.7 | -0.37 | 1956.9 | 3.75 | -11.17 | 19.8 |
| G101 | Andean | 1964.8 | 1.32 | 1890.6 | 3.78 | 39.72 | 69.3 |
| G138 | Andean | 1846.8 | 0.22 | 1826.1 | 3.25 | 6.50 | 31.0 |
| G180 | Andean | 1838.9 | 0.09 | 1789.6 | 3.13 | 2.57 | 29.5 |
| G162 | Andean | 1828.7 | 0.57 | 1814.3 | 3.20 | 16.97 | 40.5 |
| G124 | Andean | 1815.0 | -0.03 | 1805.5 | 3.19 | -0.78 | 26.0 |
| G173 | Andean | 1792.6 | 0.68 | 1780.4 | 3.07 | 20.40 | 46.0 |
| G115 | Andean | 1788.9 | 0.40 | 1765.4 | 3.04 | 11.87 | 35.5 |
| G150 | Andean | 1758.8 | 0.40 | 1750.8 | 2.98 | 12.03 | 36.5 |
| G127 | Andean | 1750.0 | -0.02 | 1733.0 | 2.94 | -0.73 | 29.0 |
| G159 | Andean | 1743.3 | 1.12 | 1694.7 | 2.90 | 33.67 | 65.3 |
| G113 | Andean | 1683.8 | 1.76 | 1548.0 | 2.35 | 52.77 | 91.5 |
| G125 | Andean | 1628.2 | -0.25 | 1590.2 | 2.49 | -7.39 | 26.8 |
| G181 | Andean | 1614.1 | 1.03 | 1585.2 | 2.44 | 31.00 | 64.5 |
| G104 | Andean | 1608.8 | 0.58 | 1601.1 | 2.49 | 17.37 | 45.3 |

GYD = grain yield, DSI = drought susceptibility index, GMP = geometric mean productivity, DTI = drought tolerance index and %GYR = percent grain yield reduction. Note: Mean Rank is the mean rank of a genotype across all the drought tolerance indices. Admixed includes genotypes that are 10 to 90% Andean or Mesoamerican according to the structure analysis results.

In general, the observed *H*^*2*^ estimates under both environments revealed that much of the observed phenotypic variation was due to the genetic component, supporting the suitability of the AMDP for GWAS studies. Grain yield was highest under NS (1016 kg/ha; *H*^2^ = 0.88), and lower under DS (715 kg/ha; *H*^*2*^ = 0.92). The SW also varied among the environments at 34.98 g/100 seeds (*H*^*2*^ = 0.97), and 31.39 g/100 seeds (*H*^*2*^ = 0.97) under NS and DS, respectively. The AMDP had a shorter duration (lower values) under DS (DPM = 90.97 days), compared to NS (DPM = 104.10 days). The same trend was observed for PH, LCC, and SC. On the other hand, LT was lower (19.75 °C) under NS environments, compared to DS environments (25.22 °C). Under DS, GYD ranged from 39.4 kg/ha to 2134 kg/ha, and exhibited a narrower range than in NS where GYD ranged from 55.0 kg/ha to 2586.0 kg/ha. The coefficient of variation (CV) ranged from 5.32 to 5.58%, 2.55 to 3.71%, 33.68 to 36.57%, 12.45 to 16.91%, 22.53 to 28.13%, 16.21 to 16.62%, 7.12 to 9.42%, and 18.90 to 35.93% for DF, DPM, GYD, SW, PH, LCC, LT, and SC, respectively. Low standard deviations (SD) were observed for LT and LCC under both environments.

Combined GYD data over two seasons across environments revealed that the highest yielding genotype was G184 (DAB91 - 2222,7 kg/ha) followed by G176 (DAB302 – 2097.5 kg/ha) and G147 (CIM-SUG07-ALS-S1-3 - 2080,1 kg/ha) (Table 3).

**Table 3.** Single nucleotide polymorphism (SNP) markers associated with agronomic and physiological traits in dry bean genotypes under drought stress conditions.

| Phenotype | SNP name | CH | SNP position on genome (bp) | MAF | Allele | Effect of allele | -log<sub>10</sub> (P) value | R<sup>2</sup> | Candidate gene |
| --- | --- | --- | --- | --- | --- | --- | --- | --- | --- |
| LT | 100106140 | 06 | 14389438 | 0.25 | A/C | 1.34 | 0.000 | 0.23 |  |
|  | 100065202 | 08 | 52504423 | 0.12 | G/A | -1.43 | 0.000 | 0.22 |  |
| DFW | 100132383 | 03 | 47240686 | 0.04 | A/G | 3.76 | 0.000 | 0.70 |  |
|  | 3381050 | 02 | 25978891 | 0.03 | C/T | 3.85 | 0.000 | 0.70 | Phvul.002G122100 |
|  | 8204238 | 10 | 42089084 | 0.06 | A/G | 2.78 | 0.000 | 0.70 |  |
|  | 8212194 | 10 | 42105474 | 0.04 | A/T | 3.54 | 0.000 | 0.69 |  |
| GYD | 3384334 | 11 | 2362591 | 0.1 | A/G | -176.67 | 0.000 | 0.44 |  |
|  | 3381526 | 11 | 2362591 | 0.09 | A/G | -174.56 | 0.000 | 0.44 |  |
|  | 3382688 | 04 | 45231105 | 0.09 | G/A | 202.90 | 0.000 | 0.43 | Phvul.004G150500 |
|  | 100061855 | 11 | 40802478 | 0.42 | T/G | 138.98 | 0.000 | 0.43 |  |
| PH | 100101387 | 05 | 34925013 | 0.03 | G/A | 17.82 | 0.000 | 0.32 |  |
|  | 8198531 | 07 | 9701750 | 0.04 | G/A | -16.03 | 0.000 | 0.31 |  |
|  | 100060987 | 01 | 42938094 | 0.04 | G/A | 15.57 | 0.000 | 0.31 | Phvul.001G172300 |
|  | 100181735 | 08 | 22152034 | 0.23 | G/A | 7.57 | 0.000 | 0.30 | Phvul.008G133100 |
|  | 3379684 | 07 | 5239949 | 0.22 | T/A | -5.62 | 0.000 | 0.30 |  |
|  | 16650827 | 07 | 51719432 | 0.09 | T/C | -8.41 | 0.000 | 0.30 |  |
|  | 3380814 | 11 | 11934462 | 0.11 | C/T | 7.15 | 0.000 | 0.30 |  |
|  | 100119463 | 08 | 6003908 | 0.03 | G/A | 15.56 | 0.000 | 0.30 | Phvul.008G065700 |
|  | 8196298 | 11 | 12212674 | 0.12 | T/G | -0.67 | 0.000 | 0.30 |  |
|  | 3379078 | 08 | 7823952 | 0.02 | C/G | 17.75 | 0.000 | 0.30 | Phvul.008G080600 |
|  | 3377272 | 11 | 9410740 | 0.16 | T/C | -6.15 | 0.000 | 0.29 |  |
|  | 3379350 | 11 | 43494132 | 0.16 | C/T | 6.35 | 0.000 | 0.29 |  |
|  | 100063156 | 10 | 7307165 | 0.44 | T/C | 4.85 | 0.000 | 0.29 |  |
|  | 3379405 | 05 | 4782514 | 0.24 | G/A | -7.80 | 0.000 | 0.29 |  |
|  | 3377900 | 11 | 9691109 | 0.14 | T/A | -6.02 | 0.000 | 0.29 |  |
| SW | 16647170 | 08 | 36620996 | 0.11 | T/C | 4.46 | 0.000 | 0.66 |  |
|  | 3383047 | 03 | 50229319 | 0.33 | G/A | -2.41 | 0.000 | 0.65 | Phvul.003G263200 |
| SC | 3380850 | 01 | 50427390 | 0.08 | T/C | -10.79 | 0.000 | 0.10 | Phvul.001G254100 |
|  | 3381030 | 02 | 33669423 | 0.04 | G/A | -10.33 | 0.000 | 0.08 |  |

CH = chromosome, DFW = days to flowering, GYD = grain yield (kg/ha), SW = 100 seed weight (g), PH = plant height (cm), LT = leaf temperature (°C), SC = stomatal conductance (mmol m<sup>-2</sup> s<sup>-1</sup>), SNP = single nucleotide polymorphism, MAF = minor allele frequency, R<sup>2</sup> = proportion of the total phenotypic variation explained by the significant SNP marker after fitting the other model effects and -log<sub>10</sub>(P) = *p* value of the association.

The drought tolerance indices for the 185 genotypes based on mean GYD are summarised in Table 3 (top 20 drought tolerant genotypes) and S2 Table (all study genotypes). The severity of DS at SVES across the 2 seasons of evaluation was moderate (DII of 0.30). Among the evaluated genotypes, G158 (SWEET WILLIAM/DAB287), G135 (DAB539), G124 (DAB487), G127 (CIM-SUG07-ALS-2), G125 (CIM-RM09-ALS-BSM-12), G138 (CZ104-72) and G184 are some of the genotypes that were less sensitive to DS based on their low DSI, %GYR and overall mean ranks across the indices. These genotypes had DSI values ranging from -0.37 (G135) to 0.16 (G184) and %GYR ranging from -11.17 (G135) to 4.69 (G184). In summary, all the top 20 drought tolerant genotypes were members of the Andean gene pool, except for G158 which is an admixture (Table 3).

### Population structure analysis

The STRUCTURE analysis results and Evanno test (Δ*K*) revealed the presence of two major sub-populations (highest Δ*K* value occurred at *K* = 2) within the AMDP of dry bean (Figs 1A, 1B). The two sub-populations correspond to the Andean and Mesoamerican domesticated gene pools. The minimum ancestry or membership coefficient to a particular cluster was 0.63 (Fig 1B and S1 Table). Most of the genotypes (90) clustered within the Mesoamerican gene pool (Fig 1B). Seventy-six genotypes clustered within the Andean gene pool (Fig 1B and S1 Table). On the other hand, 19 were Andean-Mesoamerican admixed genotypes of the two gene pools (10 to 90% Andean or Mesoamerican). The admixed genotypes included SMC16, SMC21, NUA674, NUA59-4, G75, DAB115, DAB63, DAB142, DAB477, CIM-RM02-36-1, CIM-RM09-ALS-BSM-11, CIM-RM02-134-1, Sweet William, ZABRA16575-60F22, GLP585/MLB49-89A-3, RWR2154, SAB792, NAVY LINE 22, and CIM-SUG07-ALS-S1-3 (S1 Table).

**Fig 1.**
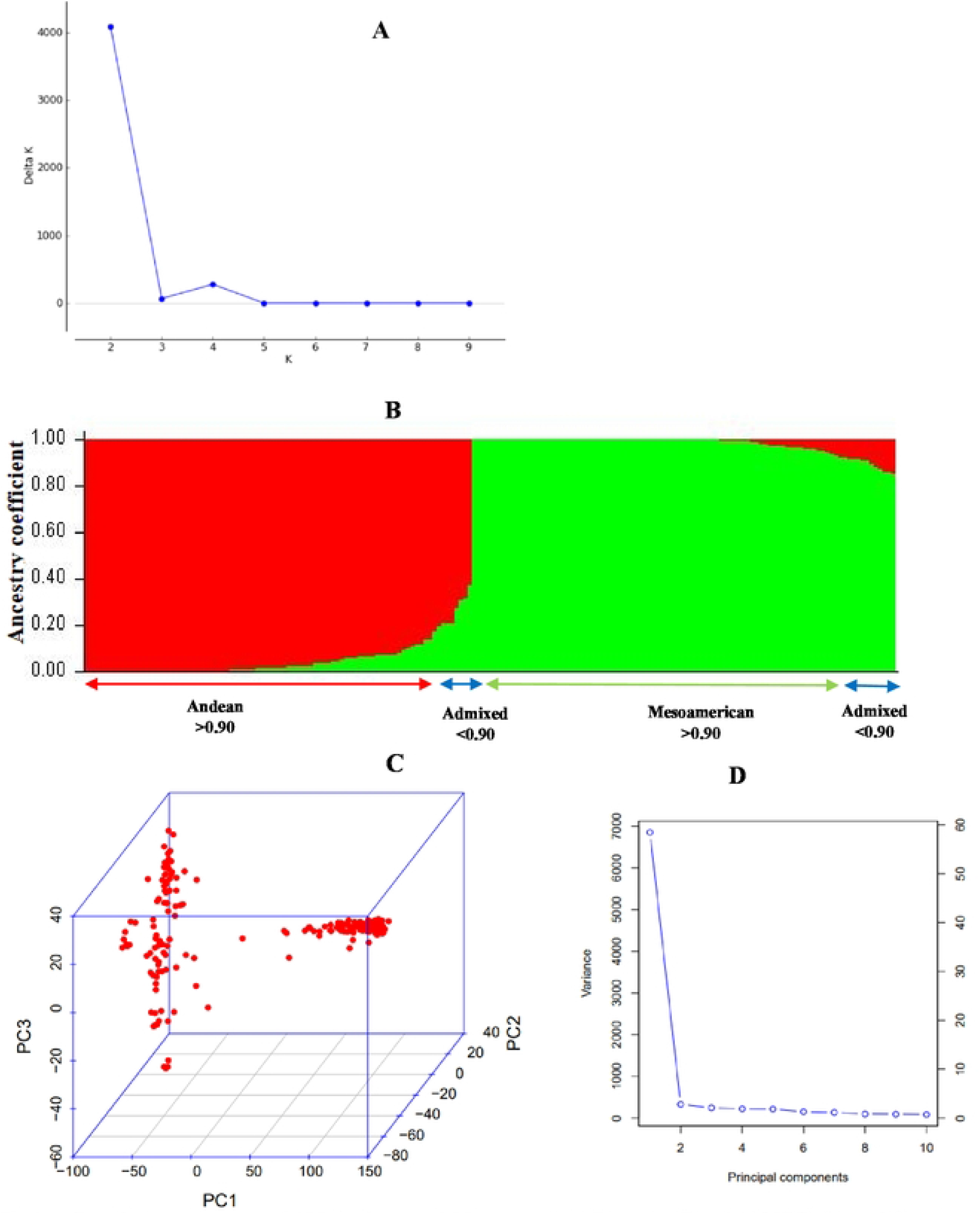
Population structure of 185 Andean and Mesoamerican Diversity Panel (AMDP) from different models. Note: A = The ΔK determined by the Evanno method showing the stratification of the 185 AMDP into two main sub-populations. The cluster with the largest ΔK (K = 2) was used to determine the number of sub-populations in the AMDP of dry bean and the existence of two-sub-populations was inferred; B = Population structure of 185 AMDP of dry bean genotypes based on 4095 SNP markers (K = 2 gives the best separation) as determined from STRUCTURE analysis. Red and green represents Andean and Mesoamerican sub-populations, respectively; C = Three dimensional principal component analysis (PCA) scatter plot illustrating the population structure of 185 AMDP of dry bean genotypes based on 9370 SNP markers; D = Screen plot showing the percentage of variation explained by the different principal components.

The genetic structure result of the AMDP was verified with the PCA based on SNP marker data and is illustrated by a 3D scatter plot (Fig 1C). The first principal component (PC) accounted for more than 55% of the observed genotypic variability in the AMDP, while the second and third PCs separately accounted for less than 5% of the overall genetic variance in the AMDP (Fig 1D). The PCA also divided the genotypes into two distinct clusters (Andean and Mesoamerican sub-populations) as were found with STRUCTURE output (Fig 1C). Further, the Andean-Mesoamerican admixed genotypes (positioned between the two groups) were isolated from the Andean and Mesoamerican sub-groups by PCA (Fig 1C).

### Analysis of marker-trait associations under drought stressed conditions

The significant MTAs and their respective statistical parameters for agronomic and physiological traits are summarised in Table 4. In this study, the threshold for significant MTA was set at p < 0.001 to reduce the risk of false MTAs. Under DS conditions, 29 significant MTAs were identified for six traits (excluding DPM and LCC) with p < 10^−03^. The associations are shown in Fig 2. The quantile-quantile (QQ) plots for the studied traits revealed that the expected and observed probability values were normally distributed (S3 Fig). The highest number of significant MTAs were observed on *P. vulgaris* (*Pv*) chromosome *Pv11* (28%), followed by *Pv8* (17%), with the least on chromosomes *Pv6* and *Pv4*, both with 3%. No significant associations for DPM and LCC were identified under DS conditions in this study. The highest number of significant MTAs were identified for PH (15), and the SNPs were distributed across six different chromosomes (*Pv1, Pv5, Pv7, Pv8, Pv10*, and *Pv11*). Additionally, the allele effect of these SNPs ranged from -16.03 cm (SNP 8198531) to 17.82 cm (SNP 100101387).

**Table 4.**
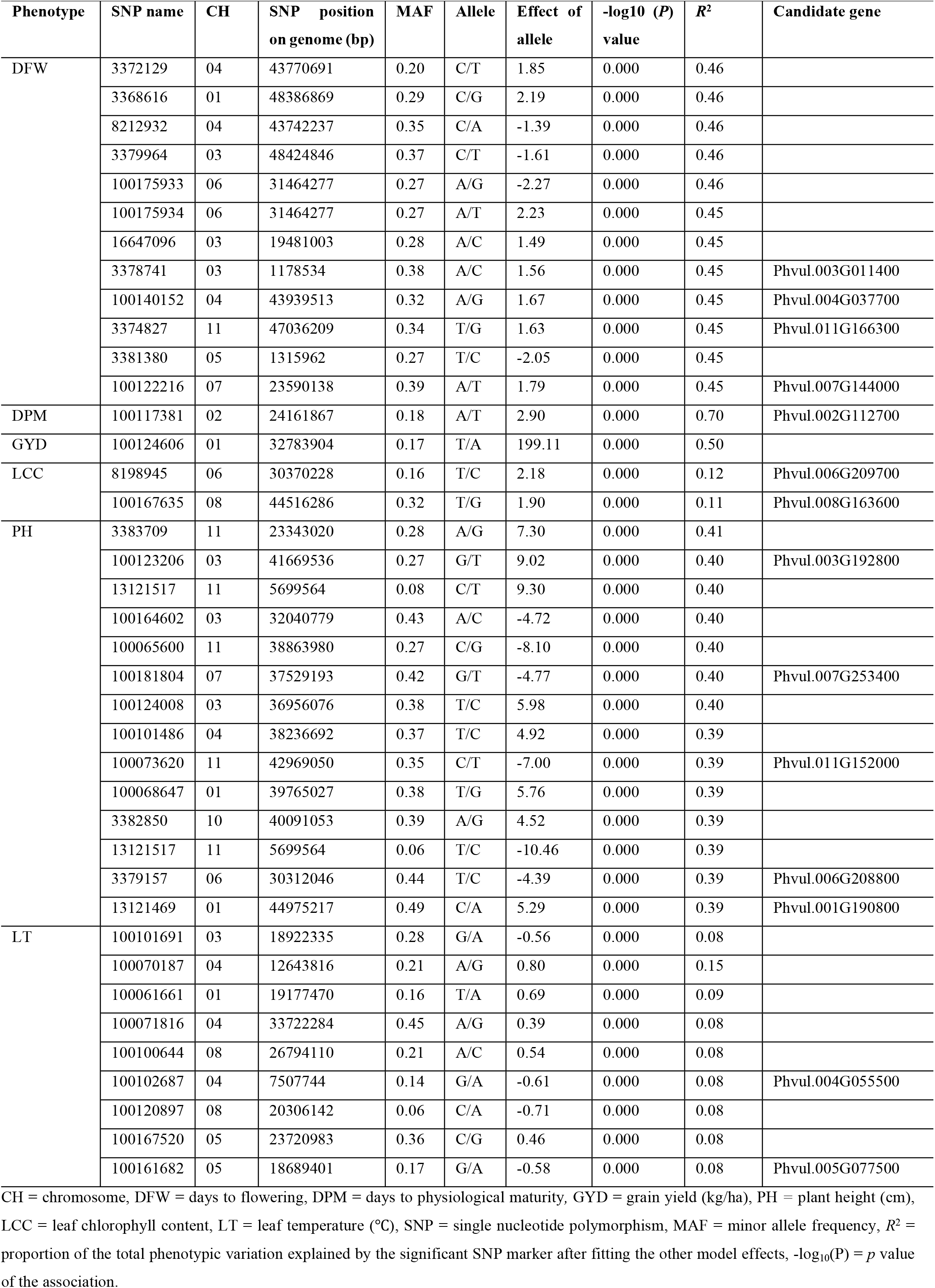
Single nucleotide polymorphism (SNP) markers associated with agronomic and physiological traits in dry bean genotypes under non-stressed conditions.

**Fig 2.**
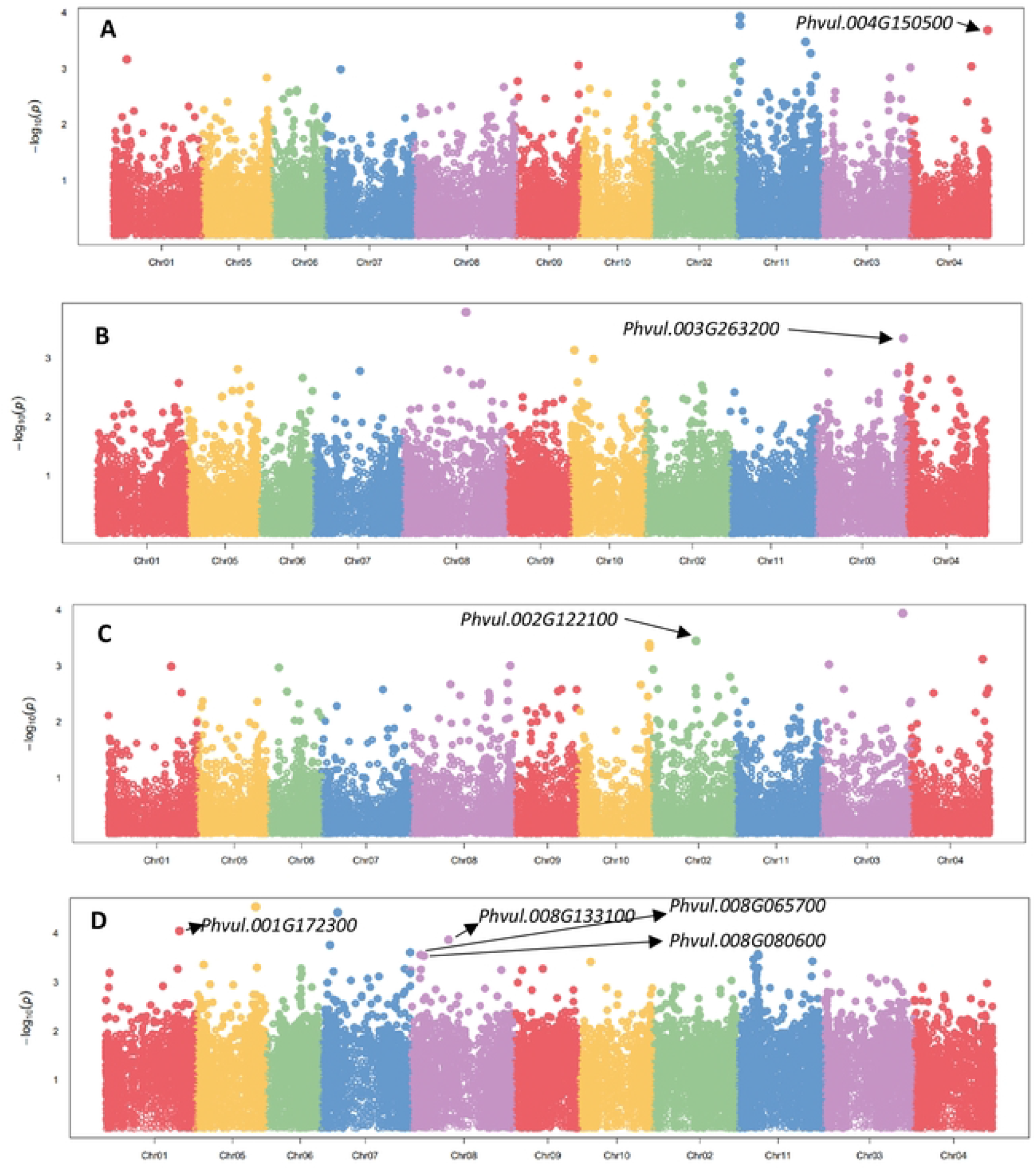
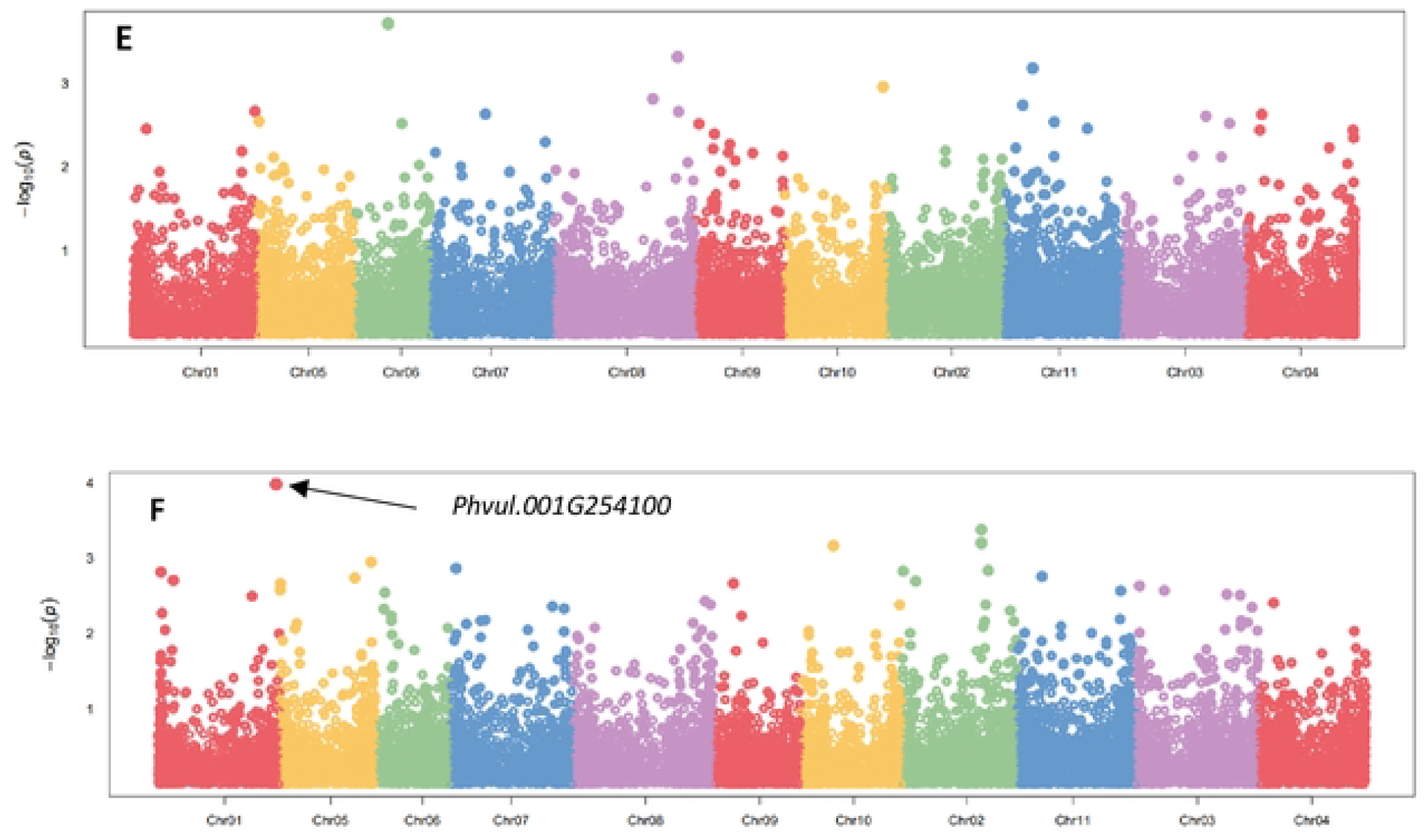
Manhattan plots indicating the significant marker-trait associations, their p-values and candidate genes for agronomic and physiological traits in 185 dry bean genotypes evaluated under drought stressed conditions. Note: A = Grain yield, B = Seed size, C = Days to 50% flowering, D = Plant height, E = Leaf temperature, F = Stomatal conductance. *Chr represents Chromosome, x-axis represents the physical map locations of the SNPs on each chromosome and the y-axis (–log base_10_ p-values) represents the degree to which a SNP is associated with a trait.

Four SNPs (SNPs 2362591, 2362591, 45231105, and 40802478) that have a significant association with GYD were also identified, and these were located on chromosomes *Pv4* and *Pv11*, with allele effect ranging from -174.56 kg/ha (SNP 3381526) to 202.90 kg/ha (SNP 3382688). Notably, 75% of the SNPs that were significantly associated with GYD were located on chromosome *Pv11*. The sum of the SNPs with a significant positive effect on GYD was 341,88 kg/ha and -351,23 kg/ha for all the SNPs with a significant negative effect on GYD (Table 4). For SW, two SNPs that were significantly associated with this trait were identified on chromosomes *Pv03* and *Pv08*, with allelic effects ranging from -2.41 g per 100 seeds (SNP 3383047) to 4.46 g per 100 seeds (SNP 16647170). Regarding physiological traits, SNPs were identified that have a significant association with LT distributed across two chromosomes (*Pv6* and *Pv8*), with allele effect ranging from -1.23°C (SNP 100065202) to 1.34°C (SNP 100106140).

Notably, two SNPs on chromosomes *Pv1* and *Pv2* were significantly associated with SC, with allele effect ranging from -10.79 mmol m^−2^ s^−1^ (SNP 3380850) to -10.33 mmol m^−2^ s^−1^ (SNP 3381030). Common regions associated with multiple traits on chromosomes were not identified under DS environments in this study. Markers explained 0.08 – 0.10, 0.22 – 0.23, 0.29 – 0.32, 0.43 – 0.44, 0.65 – 0.66 and 0.69 – 0.70 of the total phenotypic variability (*R*^*2*^) for SC, LT, PH, GYD, SW and DFW, respectively. Overall, the *R*^*2*^ varied from 0.08 (SC: SNP 3381030) to 0.70 (DFW: SNPs 100132383, 3381050 and 8204238).

### Analysis of marker-trait associations under non-stressed environments

The significant MTAs and their respective statistical parameters for agronomic and physiological traits are summarised in Table 5. Under NS conditions, 39 significant MTAs were detected for six traits (excluding SW and SC) with p < 10^−03^. The associations are shown in Fig 3.

**Fig 3.**
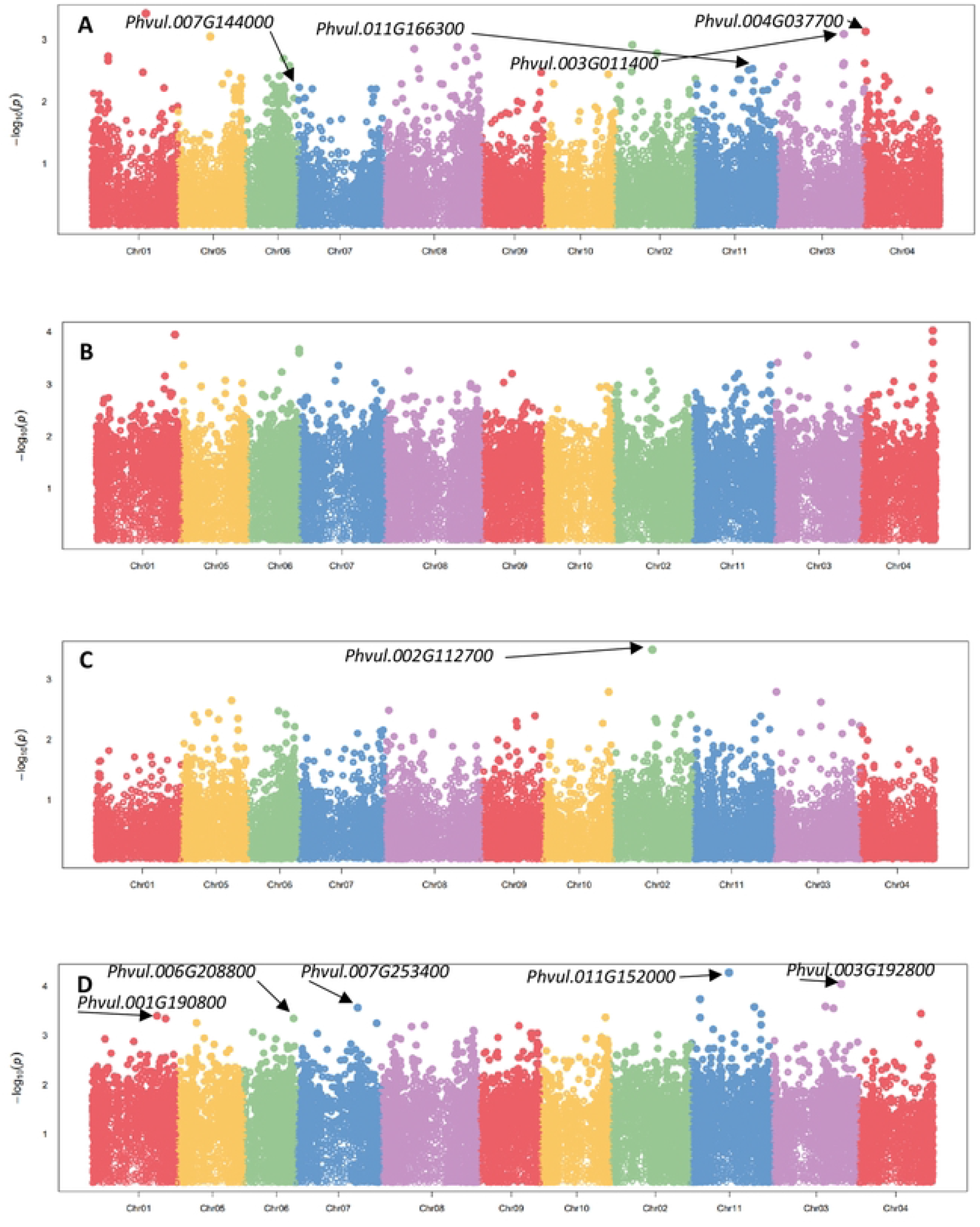
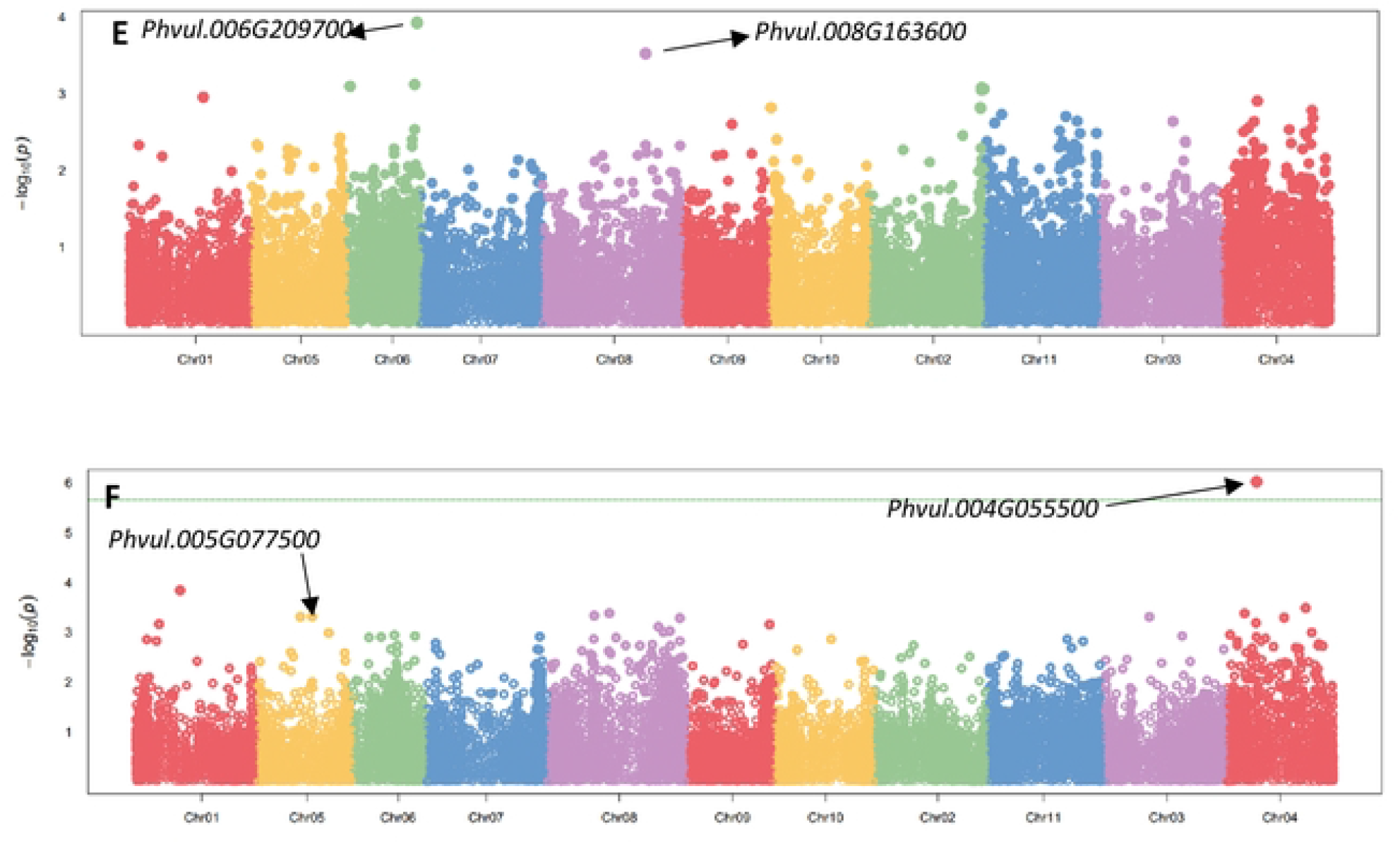
Manhattan plots showing significant marker-trait associations, their p-values and candidate genes for agronomic and physiological traits under well-watered conditions. Note: A = Days to 50% flowering, B = Grain Yield, C = Days to physiological maturity, D = Plant height, E = Leaf chlorophyll content, F = Leaf temperature. *Chr represents Chromosome, x-axis represents the physical map locations of the SNPs and the y-axis (–log base_10_ p-values) represents the degree to which a SNP is associated with a trait.

The quantile-quantile (QQ) plots for the studied traits revealed that the expected and observed probability values were normally distributed (S4 Fig). The highest number of significant MTAs were observed on *Pv11* (15%), followed by chromosomes *Pv3* and *Pv4* (both with 18%), with the least on *Pv2* and *Pv10* (both with 3%). No significant markers for SW and SC were detected under NS conditions in this study. The highest number of significant MTAs were observed on PH (14), with markers accounting for 0.39 – 0.40 of the total trait variation. Additionally, the allele effect of these SNPs ranged from -10.46 cm (SNP 13121517) to 9.30 cm (SNP 13121517). Interestingly, 38% of the markers that were significantly associated with PH were located on chromosome 11. For DFW, a total of 12 significant associations were identified, with markers explaining 0.45 – 0.46 of the observed trait variation. Additionally, the significant SNPs for DFW were located on chromosomes *Pv1, Pv3, Pv4, Pv5, Pv6, Pv7* and *Pv11*, with allele effect ranging from -2.27 days (SNP 100175933) to 2.23 days (SNP 100175934).

Notably, one SNP (SNP 100124606) on chromosome *Pv01* was significantly associated with GYD, with a large positive allelic effect of 199.11 kg/ha. In addition, this SNP had a MAF of 0.17 in the population. Regarding physiological traits, SNPs were identified that have a significant association with LCC distributed across two chromosomes (*Pv6* and *Pv8*), with positive allele effects ranging from 1.90 (SNP 100167635) to 2.18 (SNP 8198945). For LT, nine significant associations were detected, with markers accounting for 0.08 – 0.15 of the trait variation. The significant SNPs for LT were located on chromosomes *Pv1, Pv3, Pv4, Pv5* and *Pv8*, with allele effect ranging from -0.71°C (SNP 100102687) to 0.80°C (SNP 100070187). Additionally, the sum of the SNPs with a significant positive effect on LT was 2.88°C and - 2.46°C for all the SNPs with a significant negative effect. A locus (SNP 100117381) on chromosome *Pv02* explained the highest proportion of the phenotypic variation (0.70) among the studied traits and was associated with DPM. In addition, SNP 100117381 had a MAF of 0.18 in the population and a large positive effect (2.90 days) on DPM. On the other hand, nine significant SNPs for LT on chromosomes *Pv3, Pv4, Pv8* and *Pv5* explained the least proportion of the observed phenotypic variation (0.08) among the studied traits. Common regions associated with multiple traits on chromosomes were not identified under NS environments. Overall, *R*^*2*^ varied from 0.08 (LT – SNPs 100101691, 100071816, 100100644, 100102687, 100102687, 100167520 and 100161682) to 0.70 (DPM - SNP 100117381) (Table 5).

### Identification of putative candidate genes associated with significant single nucleotide polymorphism

#### Drought stressed environments

A total of eight potential candidate genes (DFW - 1; GYD - 1; PH - 4; SW - 1; SC – 1) were identified under DS environments (Table 4 and Fig 2). The candidate genes for DFW (*Phvul*.*002G122100*), SC (*Phvul*.*001G254100*), SW (*Phvul*.*003G263200*) and GYD (*Phvul*.*004G150500*) were identified on chromosomes *Pv02, Pv01, Pv03* and *Pv04*, respectively (Table 4). These genes had diverse putative functions ranging from RNA recognition motif or RNP domain functions (DFW), NADPH dehydrogenase/NADPH diaphosare activity (SW), helicase activity and CCCH zinc finger protein domain functions (SC) to Phosphoethanolamine N-methyltransferese activity (GYD), respectively. On the other hand, the candidate genes for PH were identified on chromosomes *Pv01* (*Phvul*.*001G172300*) and *Pv08* (*Phvul*.*008G133100*; *Phvul*.*008G065700*; *Phvul*.*008G080600*) (Table 4). These genes had diverse putative functions ranging from calcium transporting ATPase 1 activity, peptidyl prolyl cis trans isomerase activity, acyl-coenzyme A thiosterase activity to centrosomal protein nuf function, respectively.

#### Non-stressed environments

A total of fourteen potential candidate genes (DFW - 4; DPM - 1; LCC - 2; PH - 5; LT – 2) were identified under NS environments (Table 5 and Fig 3). The candidate genes for DFW were identified on chromosomes *Pv03* (*Phvul*.*003G011400*), *Pv04* (*Phvul*.*004G037700*), *Pv07* (*Phvul*.*007G144000*) and *Pv11* (*Phvul*.*0011G166300*), whereas the candidate gene for DPM was identified on chromosome *Pv02* (*Phvul*.*002G112700*) (Table 5). Candidate genes for DFW had diverse putative functions related to SORTING NEXIN-13, transcription factor TCP 13, U6 SNRNA-associated SM LIKE PROTEIN LSM4 and NHL domain containing protein. On the other hand, the candidate gene for DPM had a putative function related to the activity of thiol disulphide oxidoreductase. Chromosomes *Pv4* and *Pv5* harboured the two candidate genes for LT namely *Phvul*.*004G055500 and Phvul*.*005G077500*, respectively (Table 5). These genes had diverse putative functions related to the mitochondrial transcription termination factor family protein and leucine rich repeat protein associated with apoptosis in muscle tissue, respectively.

The genes *Phvul*.*006G209700* and *Phvul*.*008G163600* for LCC were identified on chromosomes *Pv06* and *Pv08*, respectively. These genes had diverse putative functions, such as premnaspirodiene oxygenase or hyoscymus muticus premnaspirodiene oxygenase activity and nucleoside triphosphate hydrolases activity, respectively. On the other hand, the candidate genes for PH were identified on chromosomes *Pv01* (*Phvul*.*001G190800*), *Pv03* (*Phvul*.*00G192800*), *Pv06* (*Phvul*.*006G208800*), *Pv07* (*Phvul*.*007G253400*), and *Pv11* (*Phvul*.*011G152000*) (Table 5). These genes also had diverse putative functions, such as f-box-like domain superfamily functions, protein NRT1 or PTR family related functions, phosphatidylserine decarboxylase activity, typa-like translation elongation factor svrs-related functions, and inactive g-type lectin s-receptor like serine or threonine protein kinase activity, respectively.

### Linkage disequilibrium analysis using significant SNP markers

The analysis of LD using SNP markers is shown in Fig 4. A high and extensive LD was observed for the common bean genome, which is expected in self-pollinated crops such as common bean. The results show that the overall LD decay across the genome of 185 common bean genotypes was 30 bp, at a cut–off of *r*^*2*^ = 0.4. Generally, there was a slow decay of LD throughout the common bean genome, and the LD extended to several mega-bases as shown in Fig 4. The population structure usually affects the extent of LD decay.

**Fig 4.**
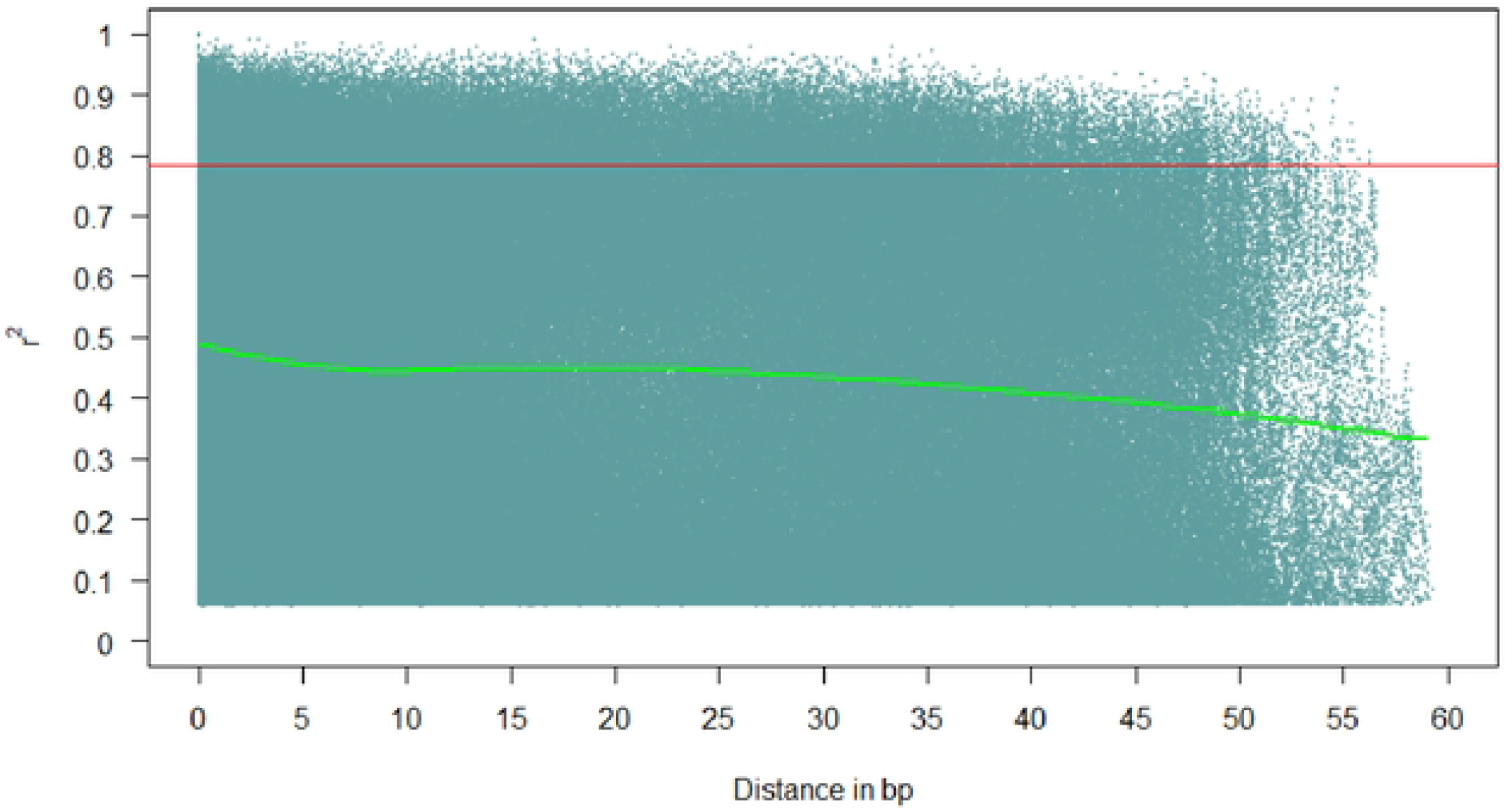
Linkage disequilibrium (LD, r^2^) decay plot in genome of dry beans based on 9370 single nucleotide polymorphisms (SNPs) in 185 diverse genotypes.

## Discussion

### Variations in agronomic and physiological traits

The low to moderate *H*^*2*^ estimates observed for SC, LT and LCC under DS and NS conditions imply that these physiological traits might be influenced by a number of genes (polygenic inheritance) and the production environment. Therefore, direct selection for SC, LT and LCC under DS and NS conditions could be a challenge to dry bean breeders. On the other hand, the high *H*^*2*^ estimates (97%) for seed size observed under DS and NS environments reflect the predominance of additive gene action (genetic control of this trait) across environments. The current findings are in agreement with Assefa et al. [80] and Hoyos-Villegas et al. [14] who reported *H*^*2*^ estimates of 77 and 93.4%, respectively under NS conditions. In this study, drought stress reduced PH, GYD, SW, DPM, LCC and SC by 12.1, 29.6, 10.3, 12.6, 28.5 and 62.0%, respectively, highlighting the detrimental effect of moisture stress under field conditions. These findings corroborate previous reports by Assefa et al. [80], Darkwa et al. [22], Assefa et al. [81], and Mathobo et al. [82] in common bean. Mathobo et al. [82] reported reductions of 48 and 39% in SC and LCC, respectively under DS conditions. Darkwa et al. [22], using navy beans, reported reductions of 10.7, 14.8, 12.7 and 26.1% in SW, PH, DPM and LCC under DS conditions. Assefa et al. [80], using navy beans, also reported reductions of 12% and 17.6% in SW and DPM, respectively under DS conditions.

Crop plants close their stomata when exposed to drought stress to minimize excessive water loss and avoid dehydration. However, the closing of stomata reduces stomatal conductance, and also affects cooling mechanisms resulting in increased leaf or canopy temperature. Therefore, in this study, drought stress increased LT by 21.6%. Drought stress also reduced GYD by 30%, close to the GYD reductions reported by Schneider et al. [83] [26%], Darkwa et al. [22] [30%] and Mutari et al. [24] [28%] in dry bean drought tolerance screening trials. Breeding for enhanced GYD under both DS and NS environments is one of the greatest challenges faced by dry bean breeders [15]. Therefore, one of the most important contribution of this study was to indicate drought tolerant genotypes (DAB91, DAB302, AFR703, CIM-SUG07-ALS-51-3, DAB487, DAB287, CIM-RM09-ALS-BSM-12 and DAB539) with consistent outstanding and stable GYD performance under both DS and NS environments. Terminal drought stress is an important factor limiting common bean productivity in the SSA region. Therefore, the identification and subsequent release of drought tolerant genotypes will positively impact on socio-economic, food and nutrition security in SSA. These genotypes could also serve as important genetic resources in drought tolerance breeding programmes to improve released cultivars. Both DAB287 and AFR703 were released in Zimbabwe as Sweet William and Gloxinia, respectively. Among the drought tolerant genotypes with superior GYD performance under water deficit conditions, most of the top 20 genotypes were of the Andean gene pool, coded as drought Andean (DAB lines) (Table 3 and S2 Table). Notably, all the DAB lines evaluated in this study were developed for improved tolerance to drought by the Alliance of Bioversity International and International Centre for Tropical Agriculture in Colombia. The current observation suggests that progress in improving drought tolerance in the Mesoamerican gene pool has been limited compared to the Andean gene pool. The current findings are in agreement with Assefa et al. [81] who reported that progress in improving drought tolerance in navy beans (Mesoamerican gene pool) worldwide has been limited compared to the other commercial classes of small seeded Mesoamerican beans.

### Population structure and Linkage disequilibrium analysis

The AMDP was delineated into two distinct major sub-populations based on the genotypes’ genetic ancestry, and this corresponded to the Andean and Mesoamerican gene pools (Figs 1B, C). This is expected considering that the domestication of dry beans on the American continent in two main centres of origin (Andean and Mesoamerican regions of America) resulted in two major and diverse gene pools [59, 84]. Cichy et al. [31, 55], Raggi et al. [74], Tigist et al. [48], Nkhata et al. [49], Ojwang et al. [71], Keller et al. [6] and Liu et al. [85] also observed two sub-populations (Andean and Mesoamerican gene pools) in their GWAS studies.

A number of the identified Andean-Mesoamerican admixed genotypes carrying genomic regions from both gene pools are released cultivars in Rwanda (RWR2154), Malawi (NUA59-4), Zimbabwe (SMC16, NUA674, and Sweet William), Eswatini (NUA674) [86–89]. Further, most of the admixed genotypes have commercial seed types, are biofortified (RWR2154, SMC16, SMC21, NUA674 and NUA59-4) and drought tolerant (Sweet William, DAB115, DAB63, DAB142 and DAB477). Singh [90], Beebe et al. [4, 20] and Beebe [84] reported that interracial hybridizations between races or sister species (*Phaseolus coccineus, Phaseolus acutifolius* and *Phaseolus dumosus*) of *Phaseolus vulgaris* have been widely used in dry bean improvement programmes when breeding for enhanced grain yield, micronutrient density and drought tolerance. For example, the biofortified admixed genotype NUA674 is a product of an inter-gene pool cross between AND277 (Andean gene pool) and G21242 (Andean-Mesoamerican inter-gene pool landrace) made at the Alliance of Bioversity International and International Centre of Tropical Agriculture (ABC) in Colombia [87]. Islam et al. [91] and Beebe [84], also reported that one of the parents to NUA674, G21242 (source of high seed iron in biofortification breeding programmes) is a product of Andean– Mesoamerican inter-gene pool hybridization, validating the current findings. Therefore, the current observation suggests that most of the admixed genotypes identified in this study resulted from deliberate breeding efforts (inter-gene pool hybridizations) to introgress genes for enhanced grain yield, drought tolerance and micronutrient density. Similar findings were reported by Hoyos-Villegas et al. [14] and Tigist et al. [48] in common bean.

The biofortified and drought tolerant admixed genotypes identified in this study may be used as a bridge to transfer favourable alleles for micronutrient density and drought tolerance into either the Andean or Mesoamerican seed types. The extent and structure of LD decay in the study germplasm usually determines the resolution of GWAS. The slow decay of LD observed in this study is expected in self-pollinating crop species, such as common bean because of the loss of recombination, which results in a homozygous genetic background. According to Vos et al. [92], recombination events in crops with a homozygous genetic background are ineffective to cause LD decay, resulting in extended (large) and slow decay of LD. The slow decay of LD, and the large extent of LD observed in this study corroborates previous reports in dry bean [32, 85].

### Marker-Trait Associations

In dry bean, it is important to enhance moisture stress tolerance by identifying genotypes with high grain yield potential under water deficit conditions, and by introgressing desirable alleles conferring drought tolerance. The mean call rate (93%) and reproducibility (100%) of the silico DArTs used in this study were consistent with previous reports [15, 49], thus demonstrating the reliability and high quality of this set of silico DArTs. A higher number of significant MTAs were detected under NS conditions, corroborating previous reports in bread wheat (*Triticum aestivum* L.) [93, 94] and dry bean [15]. The observed trend could be due to the fact that drought tolerance is a complex polygenic trait which is highly influenced by the production environment, resulting in unpredictable performance of genotypes (genotype-by-environment interaction [GEI]) under different environments (DS and NS). Even though a smaller number of significant MTA was observed under DS compared to the NS condition, novel genomic regions associated with key agronomic and physiological traits were detected under DS conditions. Notably, no significant SNPs for all the studied agronomic and physiological traits were consistent across DS and NS treatments. Similar findings were reported in wheat ([93] – plant height and spike length) and dry bean ([15] – grain yield) under DS and NS treatments. The observed trend suggests that some markers may influence the expression of phenotypic traits differently under DS and NS environments. Further, the GEI could have confounded the identification significant SNPs that are consistent across DS and NS treatments.

The highest number of significant SNPs were identified for PH. Similar findings were reported by Sukumaran et al. [95] who observed 30 significant MTAs for PH in durum wheat (*Triticum turgidum* L. ssp. *Durum*). Some of the SNPs identified in this study were located on genomic regions that had been previously reported to be harbouring genes and QTLs for the studied traits. For example, in this study, chromosomes *Pv01, Pv03, Pv04, Pv06* and *Pv07* harboured 1 SNP, 4 SNPs, 3 SNPs, 2 SNPs and 1 SNP, respectively that were significantly associated with DFW under optimal conditions. These results are consistent with Dramadri et al. [34], Nkhata et al. [49] and Keller et al. [6]. Dramadri et al. [34] identified 2 QTLs that were associated with DFW on *Pv03* under DS and NS conditions. Nkhata et al. [49] identified 2 and 5 SNPs that were significantly associated with DFW on *Pv03* and *Pv06*, respectively under NS conditions. Further, Keller et al. [6] identified 6 SNPs, 1 SNP and 1 SNP that were significantly associated with DFW on *Pv01, Pv04* and *Pv07*, respectively under optimal conditions. These findings suggest that the aforementioned QTL regions are stable across different environments and genetic backgrounds. In addition, these findings also suggest that chromosomes *Pv01, Pv03, Pv04, Pv06* and *Pv07* harbour genes for controlling flowering.

In this study, only one marker (SN 1667170) was significantly associated with SW on chromosome *Pv08* under DS conditions. These results are in accordance with Moghaddam et al. [57], and Valdisser et al. [15] who identified significant MTAs for SW on chromosome *Pv*8 under DS and NS environments, suggesting that this QTL is stable across different environments and genetic backgrounds. On the contrary, several significant MTAs for SW were previously identified under DS on chromosome *Pv01*, [52], chromosome *Pv03* [51], chromosome *Pv09* [14], and chromosomes *Pv2* to *Pv4* and *Pv6* to *Pv11* [15]. Thus, the detection of significant MTAs for SW on different chromosomes and locations indicates high genetic diversity in common bean with respect to genomic regions associated with SW under drought stress. In this study, the identified SNPs that were significantly associated with GYD under DS were located on chromosomes *Pv04* (SNP 3382688) and *Pv11* (SNP 3384334 and SNP 3381526). Similarly, Dramadri et al. [34] identified significant QTL signals for GYD and yield components on chromosomes *Pv01, Pv02, Pv03, Pv04, Pv06*, and *Pv11* under DS conditions. Oladzad et al. [96] also identified SNPs that were significantly associated with GYD, placed on chromosomes *Pv03, Pv08*, and *Pv11* under heat stress. Further, Valdisser et al. [15] found 25 QTLs that were associated with GYD on chromosomes *Pv02, Pv03, Pv04, Pv08, Pv09* and *Pv11* under NS conditions, in agreement with the current findings. These findings suggest that chromosomes *Pv04* and *Pv11* harbour genes for controlling GYD.

The identification of SNPs associated with GYD, under moisture stress, would significantly contribute to the development of molecular tools for MAS and identification of genes of interest for edition. The proportion of the total phenotypic variation (*R*^*2*^) explained by the significant SNP markers for LCC and LT was generally low (0.11 – 0.12 for LCC under NS and 8 – 15% for LT under NS). Therefore, to account for the missing variation, it might be worthwhile to complement the SNP-based GWAS by haplotype-based GWAS [97].

### Candidate genes

#### Drought stressed

The functional annotation revealed that the candidate gene for SC, *Phvul*.*001G254100* on chromosome *Pv01* encodes the CCCH zinc finger family protein which plays an important function in response of plants to biotic and abiotic stresses [98–101]. This functional gene also plays an important role in physiological and plant developmental processes [101]. Similar findings were reported in *Brassica rapa* [98], common bean [15] and Barley (*Hordeum vulgare* L.) [101]. Wang et al. [102], Seong et al. [103] and Selvaraj et al. [104] reported that several types of CCCH zinc family finger millet genes such as *O*_*s*_*C*_*3*_*H*_*10*_, *O*_*s*_*C*_*3*_*H*_*47*_, and *OsTZF*_*5*_ are involved in the regulation of tolerance to moisture stress in rice (*Oryza Sativa* L.). According to Lin et al. [105], the CCCH zinc finger family gene confers drought tolerance in plants by regulating the opening and closing of stomata. They further reiterated that genotypes that are tolerant to drought stress have abnormal and lower stomatal conductance under moisture stressed conditions. In this study, the marker SNP 3380850 for the gene *Phvul*.*001G254100* which confers tolerance to drought stress exhibited negative allelic effects (-10.79 mmol m^−2^ s^−1^) on SC.

The functional annotation revealed that the candidate gene for DFW, *Phvul*.*002G122100* on chromosome *Pv02* encodes an RNA-recognition motif protein, which plays a comprehensive biological function (critical modulators) in abiotic stress (drought, heat flooding, cold and high salinity) responding processes in plants [106]. Zhou et al. [107] observed that the RNA-recognition motif gene “OsCBP20” from rice confers abiotic stress tolerance in *escherichia coli*. Therefore, the candidate gene *Phvul*.*002G122100* identified in this study may play a protective role under DS conditions. Candidate genes such as *Phvul*.*003G263200* (*Pv08*) for SW which encodes for NADPH dehydrogenase plays an important role in mechanisms which protect plants against nitro-oxidative stresses generated by biotic and abiotic stresses such as drought, low temperature, heat, and salinity [108]. Under DS, the seed is significantly affected by oxidative damages, and oxidative damages are minimized by the activity of NADPH dehydrogenase [109].

The candidate gene for GYD, *Phvul*.*004G150500* on chromosome *Pv04*, encodes the enzyme, phosphoethanolamine N-methyltransferese in plants. This catalytic enzyme plays an important role in the response of plants to abiotic stresses such as drought and salt tolerance by catalysing the methylation of phosphoethanolamine to phosphocholine [110]. Studies conducted by Wang et al. [110] in transgenic tobacco revealed that phosphoethanolamine N-methyltransferese improved the drought tolerance of transgenic tobacco. Notably, the marker (SNP 3382688) for this candidate gene *Phvul*.*004G150500* had large positive allelic effects (202.90 kg/ha) on GYD. The candidate gene for PH, *Phvul*.*001G172300* encodes the calcium transporting ATPase, which plays an important role in growth and development processes, opening and closing of stomata, hormonal signalling, and regulation of responses to biotic and abiotic stresses in plants [111]. In summary, these results further confirmed that the identified putative potential candidate genes were associated with moisture stress tolerance of dry bean. Therefore, the putative candidate genes identified in the current AMDP under DS conditions are important genetic resources. The candidate genes could be utilized in drought tolerance breeding programmes by creating and introgressing new genetic variability into commercial cultivars.

#### Well-watered conditions

The functional annotation revealed that the candidate gene for PH “*Phvul*.*011G152000*” on chromosome *Pv11* encodes the threonine protein kinase, which is associated with enhanced tolerance to biotic and abiotic stresses in plants [15]. Similar results were reported in dry beans by Valdisser et al. [15]. In rice, kinase causes dwarfism by reducing plant height [112]. Similarly, in this study, the marker SNP 100073620 for the gene “*Phvul*.*001G152000*” exhibited negative allelic effects (-7.00 cm) on PH. According to Zhang et al. [112], kinases also has an impact on grain yield. The candidate gene *Phvul*.*004G037700* which was found on chromosome *Pv04* in association with DFW encodes transcription factor TCP_13_. The transcription factor families are strongly involved in abiotic and biotic stress responses, including zinc-finger, dehydration-responsive element-binding (DREB), and basic helix-loop-helix (bHLH) families which regulate plant growth in leaves and roots under water deficit conditions [113]. Studies conducted by Urano et al. [113] in *Arabidopsis thaliana* revealed that TCP_13_ induces changes in leaf (leaf rolling and reduced leaf growth) and root morphology (enhanced root growth). This results in enhanced tolerance to dehydration stress under osmotic stress. The candidate gene *Phvul*.*004G055500* which was found in association with LT on chromosome *Pv04* encodes mitochondrial transcription termination factor family protein. According to Kim et al. [114], the mitochondrial transcription termination factor family protein enhances thermo-tolerance in *Arabidopsis*.

## Conclusions

This study contributes many significant MTAs in common bean for agronomic and physiological traits under DS and NS environments. The present study identified a total of 68 SNPs that were significantly (p < 10^−03^) associated with key agronomic and physiological traits under DS and NS conditions. The highest number of significant MTAs were observed on chromosome *Pv11* in both environments. For the two environments (DS and NS), no common SNPs for the studied traits was detected. Overall, twenty-two potential candidate genes were identified across environments. Most of the identified genes had known biological functions related to regulating drought stress response, and growth and development under drought stress. The information generated from this study provides insights into the genetic basis of agronomic and physiological traits under DS stress and NS conditions, and lays the foundation for future validation studies of drought tolerance genes in dry bean. Thus, the significant MTAs identified in this study should be explored and validated further to estimate their effects using segregating populations and in different genetic backgrounds before utilization in gene discovery and marker-assisted breeding for drought tolerance. Further, functional characterization and the application of gene knockout to the identified putative candidate genes would further confirm their roles in regulating drought stress response, and growth and development under DS and NS conditions. More powerful statistical genetics tools such as genomic prediction models would be needed to identify minor genes that are associated with agronomic and physiological traits. The admixed genotypes identified in this study offer potential as genetic resources in drought tolerance and biofortification breeding programmes, especially within the sugar, red mottled and navy bean market classes.

## Acknowledgements

The authors would like to thank the Pan Africa Bean Research Alliance (PABRA) for the technical support received during the field and laboratory work. The authors are grateful to the Department of Research and Specialist Services (DR&SS) for providing their experimental sites for the field work. We would also like to thank PABRA for providing the germplasm.

## Author Contributions

**Conceptualization:** Bruce Mutari, Julia Sibiya, Edmore Gasura

**Data Curation:** Bruce Mutari, Admire Shayanowako, Charity Chidzanga

**Formal analysis:** Bruce Mutari, Admire Shayanowako, Charity Chidzanga

**Investigation:** Bruce Mutari

**Funding acquisition:** Bruce Mutari

**Methodology**: Bruce Mutari, Julia Sibiya, Edmore Gasura, Prince Matova

**Software:** Admire Shayanowako, Charity Chidzanga

**Resources:** Bruce Mutari

**Supervision:** Julia Sibiya, Edmore Gasura

**Validation:** Bruce Mutari, Julia Sibiya, Edmore Gasura, Admire Shayanowako, Charity Chidzanga

**Writing – original draft**: Bruce Mutari, Julia Sibiya

**Writing – review** & **editing:** Bruce Mutari, Julia Sibiya, Edmore Gasura, Admire Shayanowako, Charity Chidzanga, Prince Matova

## Supporting information

**S1 Table. List of common bean genotypes used in the study, their sources and structure membership coefficient (K2) for K = 2**.

**S2 Table. Grain yield (across environments), drought susceptibility index, geometric mean productivity, drought tolerance index and percent grain yield reduction of the 185 Andean-Mesoamerican Diversity Panel**.

**S3 Fig. Quantile –Quantile (QQ) of the p-values observed and the expected from the genome-wide association study under drought stressed conditions**. Note A = Leaf temperature, B = Days to 50% flowering, C = Grain yield, D = Plant height, E = Seed size, F = Stomatal conductance.

**S4 Fig. Quantile –Quantile (QQ) of the p-values observed and the expected from the genome-wide association study under well-watered conditions**. Note A = Days to 50% flowering, B = Days to physiological maturity, C = Grain yield, D = Leaf chlorophyll content, E = Plant height, F = Leaf temperature.

